# Diffusive mediator feedback explains the health-to-disease transition of skin inflammation

**DOI:** 10.1101/2023.05.28.542659

**Authors:** Maki Sudo, Koichi Fujimoto

## Abstract

The spatiotemporal dynamics of inflammation provide vital insights into the understanding of skin inflammation. Skin inflammation primarily depends on the regulatory feedback between pro- and anti-inflammatory mediators. Healthy skin exhibits faded erythema. In contrast, diseased skin exhibits expanding erythema with diverse patterns, clinically classified into five types: circular, annular, arcuate, gyrate, and polycyclic. Inflammatory diseases with expanding erythema are speculated to result from the overproduction of pro-inflammatory mediators. However, the mechanism by which feedback selectively drives the switch from a healthy fading erythema to each of the five types of diseased expanding erythema remains unclear. This study theoretically elucidates the imbalanced production between pro- and anti-inflammatory mediators and prospective treatment strategies for each expansion pattern. Our literature survey showed that eleven diseases exhibit some of the five expanding erythema, suggesting a common spatiotemporal regulation underlying different patterns and diseases. Accordingly, a reaction-diffusion model incorporating mediator feedback reproduced the five observed types of diseased expanding and healthy fading patterns. Importantly, the fading pattern transitioned to the arcuate, gyrate, and polycyclic patterns when the productions of anti-inflammatory and pro-inflammatory mediators were lower and higher, respectively, than in the healthy condition. Further depletion of anti-inflammatory mediators caused a circular pattern, whereas further overproduction of pro-inflammatory mediators caused an annular pattern. Mechanistically, the bistability due to stabilization of the diseased state exhibits circular and annular patterns, whereas the excitability exhibits the gyrate, polycyclic, arcuate, and fading patterns as the threshold of pro-inflammatory mediator concentration relative to the healthy state increases. These dynamic regulations of diffusive mediator feedback provide effective treatment strategies for mediator production wherein skins recover from each expanding pattern toward a fading pattern. Thus, these strategies can estimate disease severity and risk based on erythema patterns, paving the way for developing noninvasive and personalized treatments for inflammatory skin diseases.

## Introduction

Spatiotemporal dynamics provide valuable insights into variability in inflammation. Normal inflammatory response occurs only in the affected area and subsides within a short period of time, whereas chronic inflammatory response expands to adjacent healthy tissue and persists for months or years [Miyake and Kaisho 2014]. Chronic inflammation is primarily attributed to an imbalance between pro- and anti-inflammatory mediators [Zhang and An 2007, Coondoo 2011, Sabat et al. 2019]. Hence, the prevention and treatment of chronic inflammation require to elucidate the mechanisms of the imbalance involved.

The possibility for direct observation makes the skin an ideal system for studying the spatiotemporal dynamics of inflammation. Skin inflammation typically manifests as redness on the skin surface and is medically referred to as erythema [Shimizu 2017]. Erythema appears when pro-inflammatory mediators (e.g., tumor necrosis factor-alpha and interleukin [IL]-1) induce vasodilation and hyperemia in the dermis (Fig. 1A). The production of pro-inflammatory mediators is influenced by characteristics of the skin, such as the skin barrier and microbiome [Bäsler and Brandner 2017, Meisel et al. 2018]. Pro-inflammatory mediators induce the production of anti-inflammatory mediators (e.g., IL-4, IL-10, and IL-13), which reduce the production of the pro-inflammatory mediators as a regulatory feedback mechanism [Opal and DePalo 2000, Zhang and An 2007]. In addition to negative feedback, pro- and anti-inflammatory mediators induce their own production via positive feedback [Nestle et al. 2009, Zhang and An 2007]. Experimental studies have revealed that dysregulation of feedback causes the overproduction of pro-inflammatory mediators and the transition from normal to chronic inflammation [Zhang and An 2007, Coondoo 2011, Sabat et al. 2019]. Normal inflammation in healthy skin appears as fading erythema, where redness decreases and eventually disappears [Tsuji 2002]. Fading erythema includes a linear pattern reflecting the affected areas in contact with, for instance, harmful animal tentacles or plant branches and a reticular pattern reflecting the capillary structure (Fig. 1B, C) [Tsuji 2002]. Erythema patterns in diseased skin differ from those in healthy skin: chronic inflammation in diseased skin appears as expanding erythema with circular, annular, polycyclic, arcuate, or gyrate patterns (Fig. 1D–H) [Willis 1978]. Erythema expands for hours or days, with multiple expanding erythema leading to fusion [Willis 1978, Shimizu 2017]. Erythema patterns provide the first clue for the diagnosis and treatment of inflammatory skin diseases regulated by mediator feedback.

**Fig. 1.**
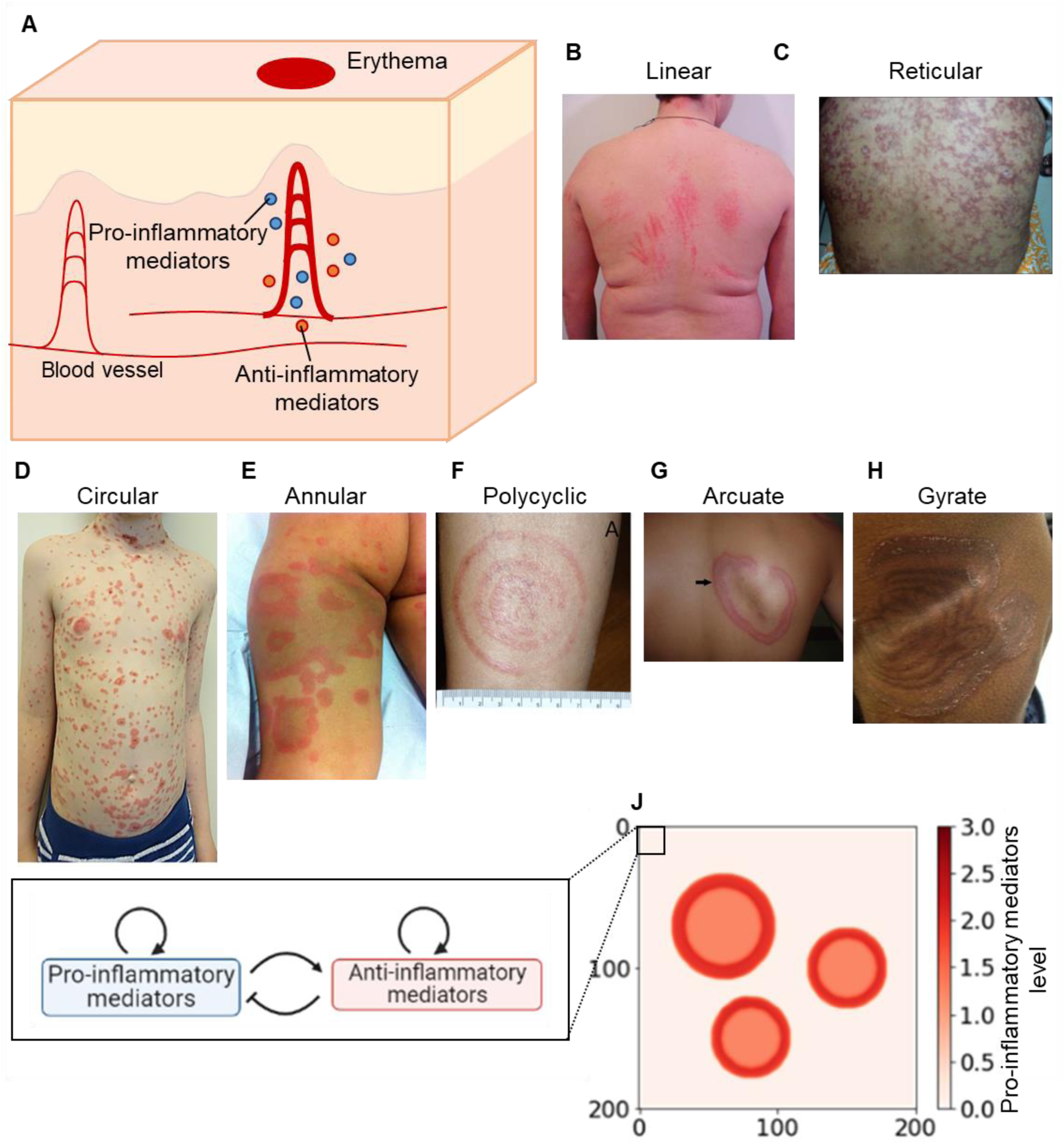
Erythema pattern and modeling of erythema development. **(A)** Process of the inflammatory response for erythema development. Upon stimulation, keratinocytes and resident immune cells secrete pro-inflammatory mediators that induce the production of pro- and anti-inflammatory mediators. Pro-inflammatory mediators dilate local blood vessels. The dilation appears as redness on the skin surface, developing erythema. **(B–H)** Photographs of erythema with linear [Adriano *et al*. 2013] (B), reticular [Naveen *et al*. 2014] (C), circular [Reichel et al. 2020] (D), annular [Abarzúa et al. 2016] (E), polycyclic [Schotthoefer et al. 2022] (F), arcuate [Jalil et al. 2020](G), or gyrate patterns [Matta et al. 2020] (H)**. (I)** A model for regulatory feedback between pro- and anti-inflammatory mediators. **(J)** A representation of simulation in the skin. The skin surface is partitioned into square regions. Erythema is initiated by keratinocytes and immune cells in the skin through secreting pro-inflammatory mediators. The area of microinflammation with a high concentration of pro-inflammatory mediators is considered as a ‘‘seed’’ region, and its projection to the surface is colored in red.

Dermatologists have reported numerous clinical findings on inflammatory skin diseases to identify appropriate treatment strategies. Clinical reports typically show the same erythema pattern in multiple diseases. For example, the annular pattern is common in erythema migrans, erythema multiforme, lichen planus, pityriasis rosea, psoriasis, tinea corporis, and urticaria [Trayes et al. 2018]. Furthermore, a clinical report comparing patients with Lyme disease revealed multiple patterns in the same disease; most New York patients developed a polycyclic pattern, whereas most Missouri patients developed an annular pattern [Wormser et al. 2005]. Moreover, three months after treatment, some New York patients remained fatigued or had joint pain, while Missouri patients did not have any of these prognostic symptoms, suggesting a correlation between erythema patterns and treatment efficacy. As these inflammatory diseases primarily result from the overproduction of pro-inflammatory mediators, mediator production can affect the development of the expanding pattern observed in different diseases. Thus, elucidating which alterations in mediator production result in specific expanding erythema patterns across diseases, enables the estimation of fundamental treatment strategies.

Mathematical modeling has recently attracted attention for predicting treatment strategies for inflammatory skin diseases. A mathematical model incorporating regulatory feedback between pro- and anti-inflammatory mediators predicts the temporal dynamics of normal and chronic inflammation [Valeyev et al. 2010]. The model characterized normal inflammation as a system with one stable steady state, where mediator concentrations transiently increased upon stimulation and subsequently returned to their original levels, showing excitability. Alternatively, chronic inflammation is characterized as a system with additional steady states with persistently high or oscillating mediator concentrations. Although the model predicted a different number of steady states underlying the temporal dynamics between normal and chronic inflammation, the absence of mediator diffusion failed to account for spatial dynamics.

Mathematical models incorporating diffusion, referred to as reaction-diffusion models, have studied the spatial dynamics of erythema patterns [Segel et al. 1992, Gilmore and Landman 2005, Penner et al. 2012, Vig and Wolgemuth 2014, Ringham et al. 2019, Seirin-Lee et al. 2020]. A reaction-diffusion model for erythema gyratum repens (EGR) suggested that the gyrate pattern characteristic of the disease is formed in the presence of excitability, where perturbations induce a transient response that returns to a stable steady state [Gilmore and Landman 2005]. Other reaction-diffusion models for psoriasis and urticaria have shown that positive and negative feedback of pro-inflammatory mediators plays a major role in generating several expanding patterns, including circular, annular, arcuate, and gyrate patterns [Ringham et al. 2019, Seirin-Lee et al. 2020]. These two models suggest that different expansion patterns within a single disease arise from alterations in mediator production due to slight differences in regulatory feedback strength. The psoriasis model also showed that the patterns faded after treatment by increasing the degradation rate of pro-inflammatory mediators. These studies focused on pro-inflammatory mediators rather than anti-inflammatory mediators. Another reaction-diffusion model incorporating pro- and anti-inflammatory mediators and chemotactic cells reproduced an expanding circular pattern [Penner et al. 2012]. Although the overproduction of pro-inflammatory mediators are thought to cause expanding erythema in many modeling studies of these inflammatory diseases [Segel et al. 1992, Gilmore and Landman 2005, Penner et al. 2012, Vig and Wolgemuth 2014, Ringham et al. 2019, Seirin-Lee et al. 2020, Sudo and Fujimoto 2022], the mechanism by which overproduction through the feedback selectively drives the switch from a healthy state with fading erythema to a disease state with each of the five types of expanding erythema remains unclear.

Elucidating this mechanism requires the development of a reaction-diffusion model for fading patterns in healthy skin. Improving the model to reproduce both fading and expanding erythema will provide better understanding of the healthy-to-disease transition and suggest non-invasive treatment strategies. Moreover, developing a model that comprehensively reproduces all five types of expanding patterns in a disease-independent manner enables us to infer how the direction and severity of the mediator imbalance affects the clinical erythema pattern.

This study aimed to theoretically elucidate how the imbalance in the production of pro- and anti-inflammatory mediators causes each expanding pattern in multiple diseases and how to restore balance and return to the fading pattern. To this end, we examined whether the expanding patterns in multiple diseases result from the reaction-diffusion system with regulatory feedback (Fig. 1A, I and J). Using the reaction-diffusion model, we explored the conditions of appearance and effective treatment strategies for each expanding pattern.

## Results

### Erythema patterns observed in eleven diseases

Previous disease-specific models have focused on multiple expanding patterns within a single disease, whereas few models have focused on studies reporting the same type of expanding pattern across different diseases [Trayes et al. 2018]. Thus, we comprehensively examined the correspondence between diseases and expanding pattern types in terms of how many diseases commonly exhibit each pattern type and the number of types each disease exhibits. The collected photographs of erythema were categorized into the following five types based on the definitions of patterns published by the International League of Dermatological Societies [Nast et al. 2016]; circular pattern characterized by a uniformly colored round pattern, annular pattern surrounded by a single ring, polycyclic pattern surrounded by multiple rings, arcuate pattern with a segmented ring, and gyrate pattern resembling wood-grain. We first examined the number of diseases that exhibited the same pattern type. Circular, annular, polycyclic, arcuate, and gyrate patterns were found in 7, 5, 4, 7, and 4 diseases, respectively (Table 1), indicating that each of the five expanding patterns corresponded to multiple diseases. We examined the number of pattern types that appeared within a single disease. Consequently, eight diseases exhibited multiple pattern types across patients. For example, psoriasis exhibited all five pattern types and lupus erythematosus exhibited four (Table 1). The most frequently observed pair of pattern types in the same disease were annular and arcuate (four diseases: psoriasis, lupus erythematosus, bullous pemphigoid, and annular erythema), whereas the least frequent were circular and gyrate (one disease: psoriasis). These results indicated that each disease corresponded to multiple pattern types. Taken together, the correspondence between patterns and diseases is many-to-many rather than one-to-one, suggesting a unified spatiotemporal regulatory mechanism across diseases to form the five types of expanding patterns.

**Table 1.** Erythema patterns observed in eleven diseases. References for each case are listed in Table S1.

| Disease name | Circular | Annular | Polycyclic | Arcuate | Gyrate |
| --- | --- | --- | --- | --- | --- |
| Psoriasis | 2 | 2 | 1 | 1 | 1 |
| Lupus erythematosus | N/A | 6 | 1 | 3 | 4 |
| Bullous pemphigoid | 2 | 3 | N/A | 3 | N/A |
| Lyme disease | 2 | 2 | 4 | N/A | N/A |
| Erythema multiforme | 3 | N/A | 8 | 1 | N/A |
| Lymphoma | 2 | N/A | N/A | 2 | N/A |
| Annular erythema | N/A | 4 | N/A | 2 | N/A |
| Sjögren syndrome | N/A | N/A | N/A | 2 | 1 |
| Sweet's syndrome | 2 | N/A | N/A | N/A | N/A |
| Nummular eczema | 2 | N/A | N/A | N/A | N/A |
| Erythema gyratum repens | N/A | N/A | N/A | N/A | 1 |
| Reported number of diseases | 7 | 5 | 4 | 7 | 4 |

### The reaction-diffusion model reproduced the fading patterns

Whether mediator production via feedback can generate and control the fading pattern, which remains uninvestigated in the reaction-diffusion models, was then examined (Fig. 1A, I and J; Eq. 2 in Methods). Given the local stimulation reflecting the shape of animal tentacles or capillary structure [Tsuji 2002, Adriano et al. 2013, Naveen *et al*. 2014], the present model reproduced a fading linear or reticular pattern, respectively (Fig. 2A and B). With circular stimulation, the inflamed area decreased in redness without changing the diameter, and the interior of the inflamed area cleared first and eventually disappeared (Fig. 2C). This result resembles the clinical situation of a fading circular pattern [Ringham et al. 2019]. During the appearance of fading patterns, mediator levels transiently increased and then decreased to their original levels (S1 Fig. A), which is consistent with the excitatory time course of the normal inflammation model without mediator diffusion [Valeyev et al. 2010]. We further analyzed the parameters that controlled fading speed. The smaller the anti-inflammatory mediator’s basal secretion rate (*p_i_*), the slower the inflamed area disappeared (Fig. 2D). Similar results were obtained when the production of pro-inflammatory mediators was high. These results demonstrate that regulatory feedback can generate a fading pattern in synergy with diffusion and control the fading speed.

**Fig. 2.**
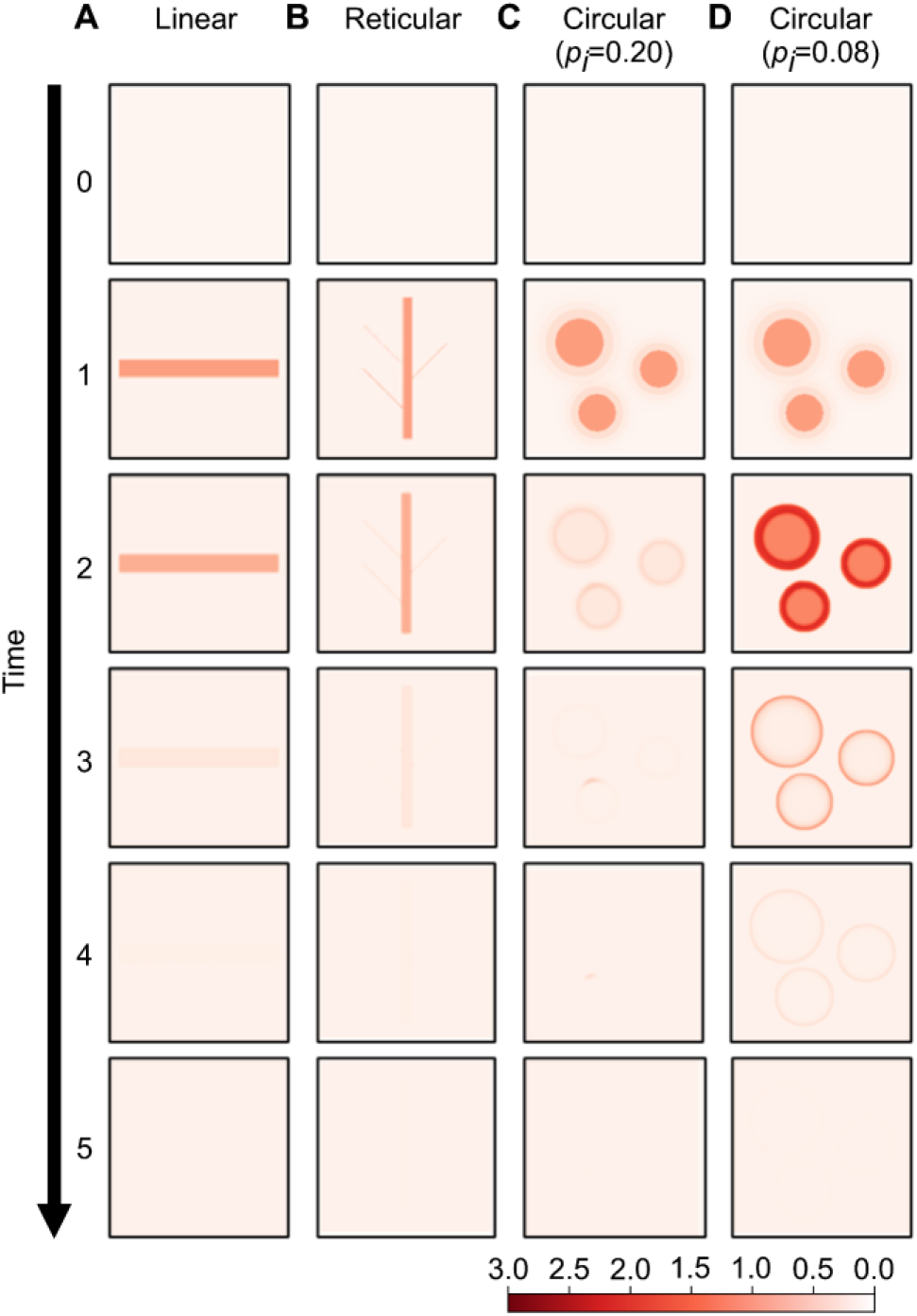
Simulated time courses of the healthy fading patterns. Spatiotemporal evolution of pro-inflammatory mediator levels (*a*; inset at the left) upon initial stimulation in linear **(A)**, reticular **(B)**, and circular areas **(C** and **D)**. The parameter values for these simulations are listed in Table S2(A).

### The reaction-diffusion model also reproduced diverse expanding patterns

We examined whether any alteration in the model parameters (Table 2) could generate five expanding patterns. The model (Eq. 2) showed that the inflamed area induced by transient local stimulation (Fig. 3, time = 1) expanded centrifugally over time (Fig. 3, time = 2–5). The inflamed area expanded with circular, annular, polycyclic, arcuate, or gyrate patterns, depending on the parameter values, such as the degradation rate of the pro-inflammatory mediator (*r_a_*) or the anti-inflammatory mediator’s basal secretion rate (*p_i_*). The circular pattern appeared as round areas with a uniform concentration of pro-inflammatory mediators above a threshold (Fig. 3A), accounting for the uniformly colored round pattern in diseased skin [Nast et al. 2016]. The annular pattern showed areas with low pro-inflammatory mediator concentrations surrounded by a single boundary ring with higher concentrations (Fig. 3B), accounting for the inflamed areas surrounded by a single ring in diseased skin [Nast et al. 2016]. The polycyclic and arcuate patterns showed double concentric rings (Fig. 3C) and segmented rings (Fig. 3D), respectively. The gyrate pattern exhibited “C”-shaped double spirals resembling wood grains (Fig. 3E). Moreover, the multiple expanding areas fused (Fig. 3, time = 2–5), consistent with the clinical situation of expanding erythema [Shimizu 2017]. Therefore, these simulated spatial patterns of pro-inflammatory mediators corresponded to each of the five types of expanding patterns in the clinical observations (Fig. 1D–H).

**Fig. 3.**
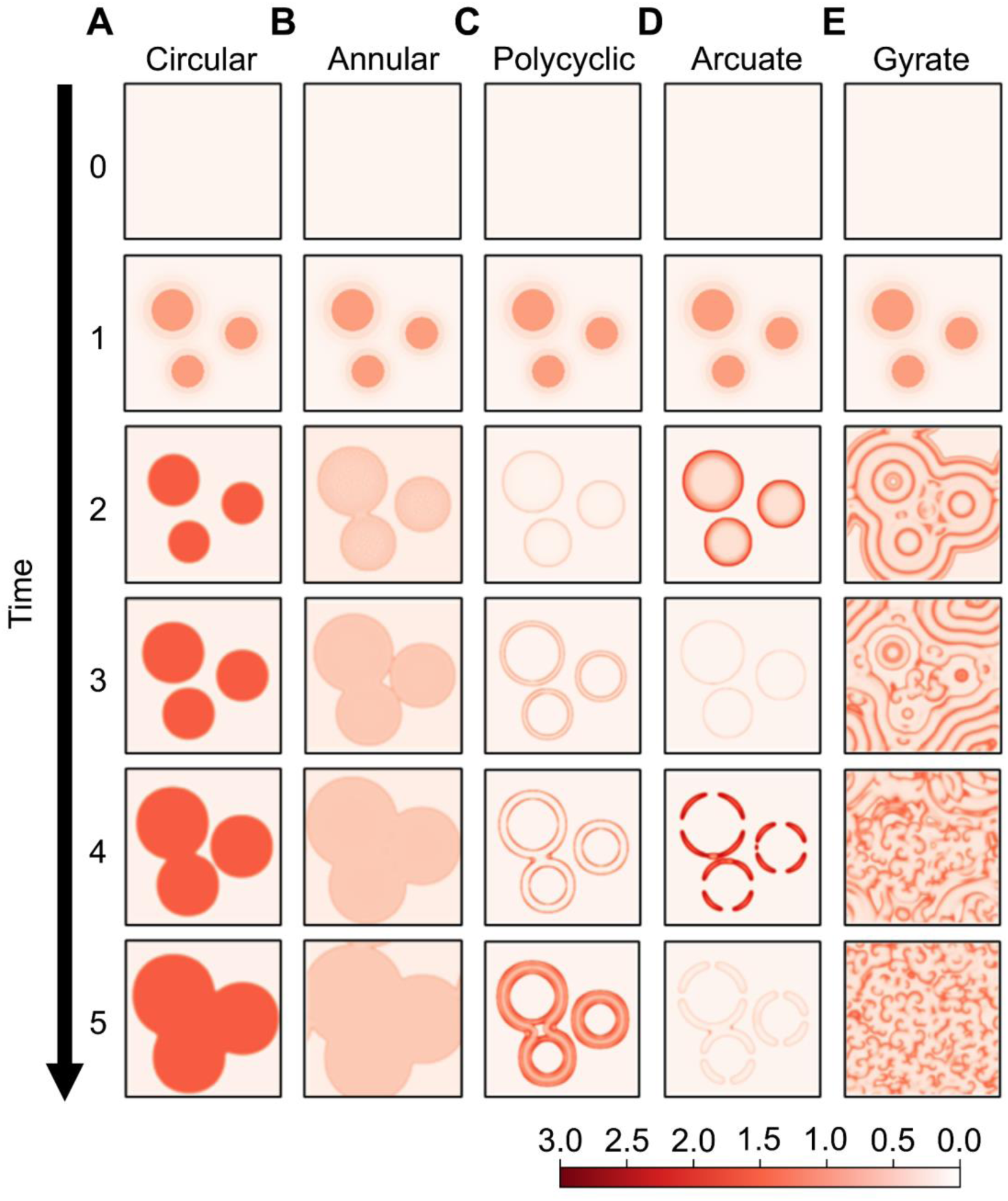
Simulated time courses of the five types of expanding patterns. The development of the expanding patterns. The initial stage of the inflamed area (row 1) consisted of three seed areas. Later forms of the disease (rows 2–5) correspond to circular **(A)**, annular **(B)**, polycyclic **(C)**, arcuate **(D),** or gyrate patterns **(E)**. The parameter values for these simulations are listed in Table S2(B).

**Table 2.**
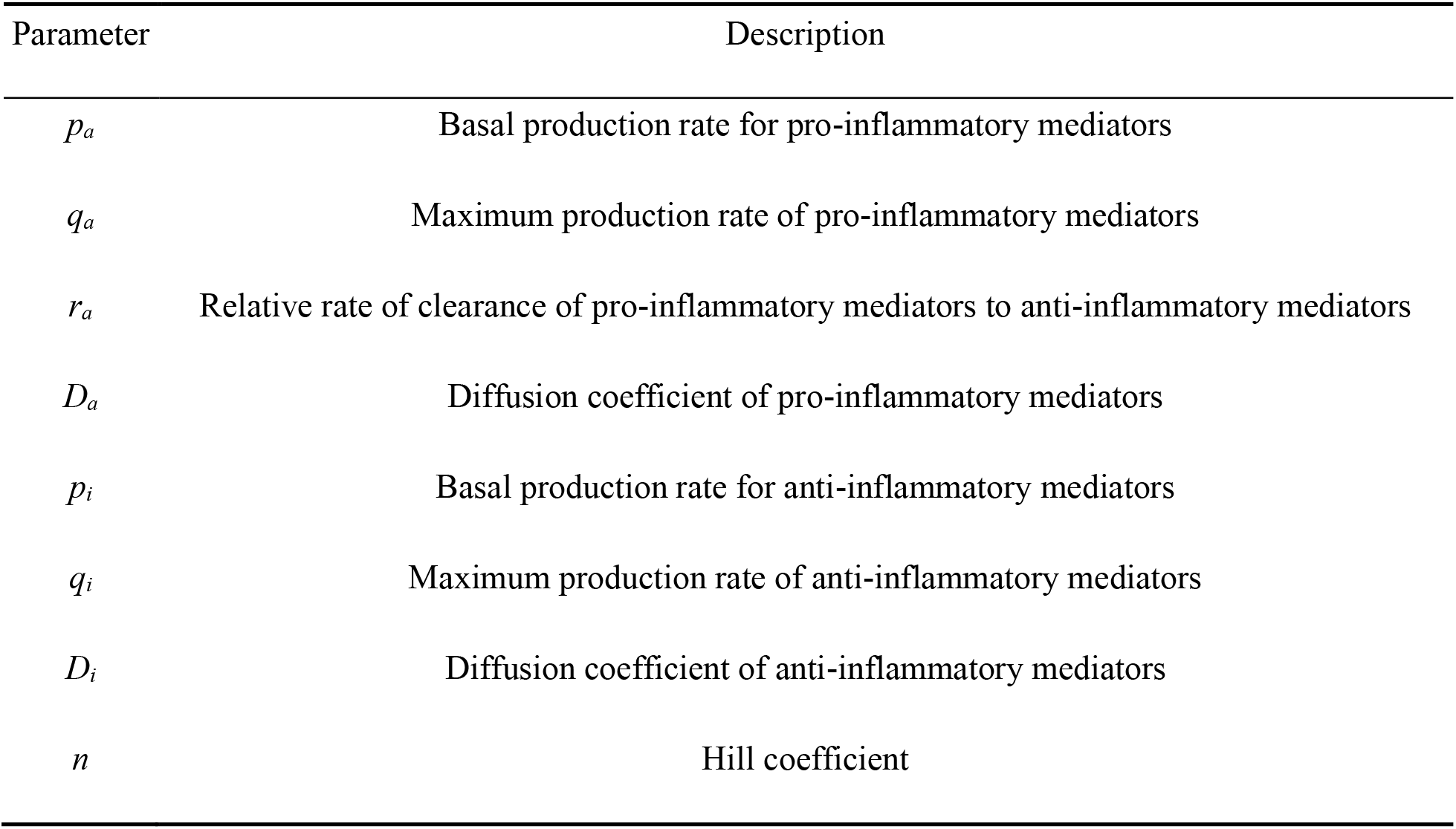
The parameters in the system and their interpretations.

| Parameter | Description |
| --- | --- |
| $p_a$ | Basal production rate for pro-inflammatory mediators |
| $q_a$ | Maximum production rate of pro-inflammatory mediators |
| $r_a$ | Relative rate of clearance of pro-inflammatory mediators to anti-inflammatory mediators |
| $D_a$ | Diffusion coefficient of pro-inflammatory mediators |
| $p_i$ | Basal production rate for anti-inflammatory mediators |
| $q_i$ | Maximum production rate of anti-inflammatory mediators |
| $D_i$ | Diffusion coefficient of anti-inflammatory mediators |
| $n$ | Hill coefficient |

The inflammatory time course was further analyzed for each expansion pattern. When a circular pattern appeared, pro-inflammatory mediators maintained a persistently high concentration and failed to return to their original level (S1 Fig. B). When the annular pattern appeared, the mediator levels transiently increased and then decreased but did not return to their original levels (S1 Fig. C). These temporal dynamics are consistent with those of chronic inflammation [Valeyev et al. 2010]. In the case of the polycyclic, arcuate, and gyrate patterns, mediator concentrations transiently increased and then returned to their original levels, indicating excitability (S1 Fig. D–F). These results support the presence of excitability in the development of gyrate patterns in EGR [Gilmore and Landman 2005]. These results indicate that the alteration of the model parameters from the fading pattern can generate five types of expanding patterns.

### Transition to expanding patterns by alteration in the production of pro- and anti-inflammatory mediators

To identify how the direction and severity of mediator production imbalance affects the pattern in the clinical spectrum of expanding patterns, we investigated the parameters (Table 2) affecting the transition from the fading pattern to each expanding pattern. First, increasing the pro-inflammatory mediator’s production rate (*q_a_*) from the parameter set of the fading pattern generated arcuate, gyrate, or polycyclic patterns (Fig. 4A). A further increase in the pro-inflammatory mediator’s production rate (*q_a_*) brought about an annular pattern (Fig. 4A). Conversely, with a decreasing basal secretion rate of the anti-inflammatory mediator (*p_i_*) from the parameter set of the fading pattern, arcuate, gyrate, polycyclic, and circular patterns appeared sequentially (Fig. 4A). These transitions from the fading pattern to all five types of expanding patterns depending on *q_a_* or *p_i_* prompted us to hypothesize that increasing pro-inflammatory or decreasing anti-inflammatory mediator concentration can cause the transition from the fading pattern to transient expanding patterns (arcuate, gyrate, and polycyclic) and ultimately to chronic expanding patterns (annular and circular).

**Fig. 4.**
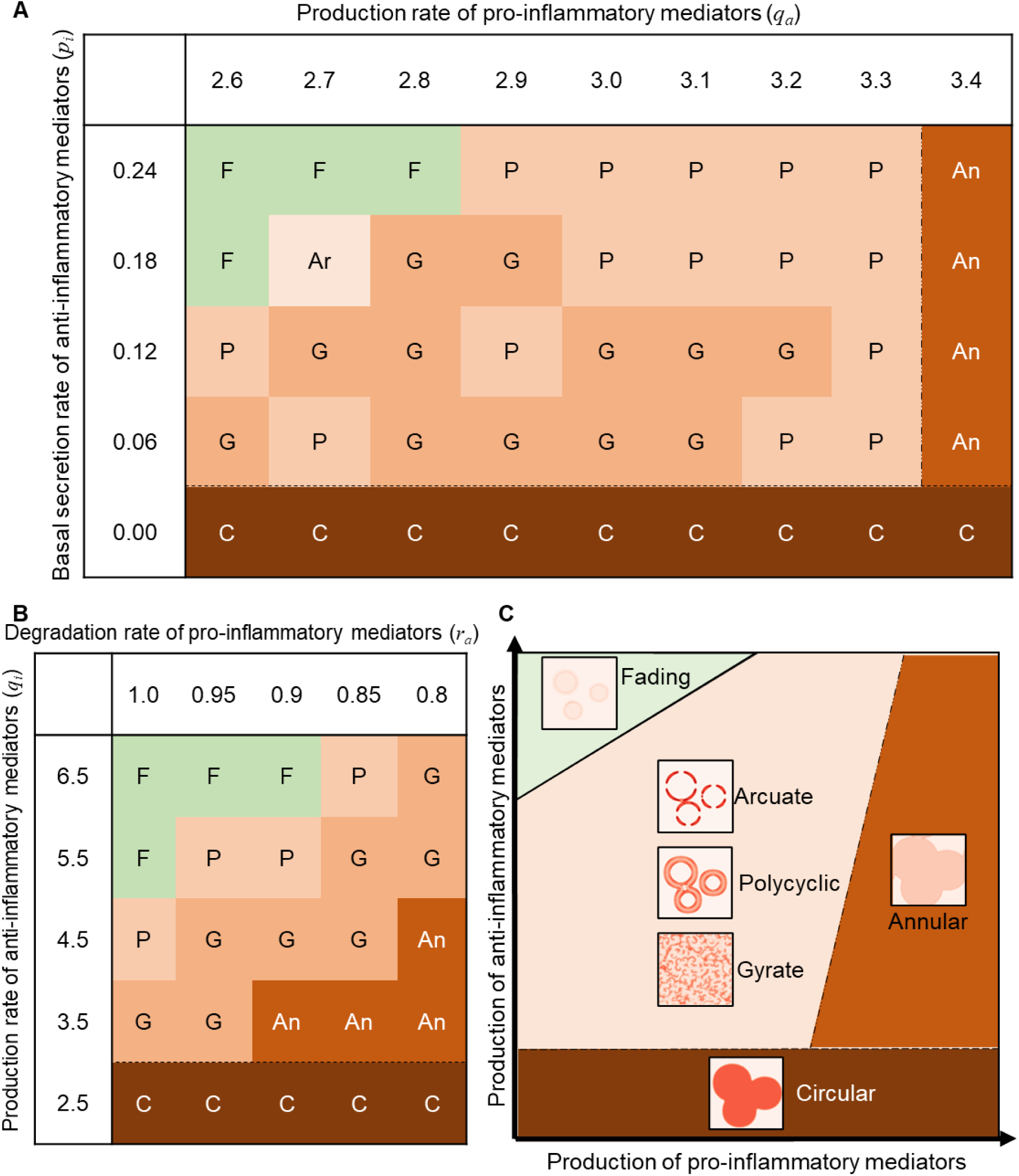
Pattern selection in the parameter space of pro- and anti-inflammatory mediator productions. Fading (F), arcuate (Ar), polycyclic (P), gyrate (G), annular (An) and circular (C) patterns emerged as the steady state (Eq. 2) at the parameter values of *q_a_* and *p_i_* **(A)**, *r_a_* and *q_i_* **(B)**. *p_a_* = 0.05, *r_a_* = 0.8, *q_i_* = 6.0 for (A) and *p_a_* = 0.05, *q_a_* = 3.0, *p_i_* = 0.12 for (B). In all the simulations, *D_a_* = *D_i_* = 0.3. **(C)** Summary for all the analyzed parameter space regarding the mediator production (see also S2 Fig. and S3 Fig.), indicating the characteristic imbalance by each expanding pattern.

To test this hypothesis, we comprehensively investigated pattern transitions with alterations in each of the parameters affecting mediator concentration. Decreasing the degradation rate (*r_a_*) of the pro-inflammatory mediator and production rate of anti-inflammatory mediators (*q_i_*) from the fading pattern parameter set consistently led to polycyclic, gyrate, and finally circular patterns (Fig. 4B), supporting this hypothesis. The results from various combinations of parameters identified the parameter regions for each expanding pattern in the clinical spectrum, ranging from transient to chronic expanding patterns (Fig. 4A, B, S2 Fig., and S3 Fig.). The transient expanding pattern, including arcuate, gyrate, and polycyclic patterns, emerged under lower production of anti-inflammatory mediators and higher production of pro-inflammatory mediators compared to the fading pattern. Excessive imbalance resulted in a chronic expansion pattern; the annular pattern appeared under the overproduction of pro-inflammatory mediators, whereas the circular pattern appeared under the depletion of anti-inflammatory mediators. Generally, alterations in all parameters of feedback in the model caused an imbalance in mediator production, resulting in transient and eventually chronic expanding patterns.

These results indicate the transition from each diseased expanding pattern to a healthy fading pattern. Specifically, the annular and circular patterns shifted to the fading pattern by reducing the production of pro-inflammatory mediators and increasing the production of anti-inflammatory mediators, respectively. Overall, these results showed that the two-dimensional space representing pro- and anti-inflammatory mediator production describes the clinical spectrum from the five types of expanding patterns in diseased skin to the fading pattern in healthy skin (Fig. 4C).

### The stability of the healthy and inflamed states determines the expanding or fading patterns

To identify the dynamic properties underlying the differences between the fading pattern and each of the expanding patterns, the number of stable states was analyzed. These states were predicted as the temporal properties between normal and chronic inflammation, which are regulated by excitability and bistability, respectively [Valeyev et al. 2010], but remain unexamined regarding the spatial patterns. In the parameter set for the circular pattern, the regulatory feedback between pro- and anti-inflammatory mediators resulted in the bistability: two steady states are stable, given by low and high concentrations corresponding to the healthy (*S_H_* in Fig. 5A) and inflamed (*S_I_* in Fig. 5A) states, respectively, whereas there is an unstable stable state, corresponding to a threshold concentration (*S_T_* in Fig. 5A).

**Fig. 5.**
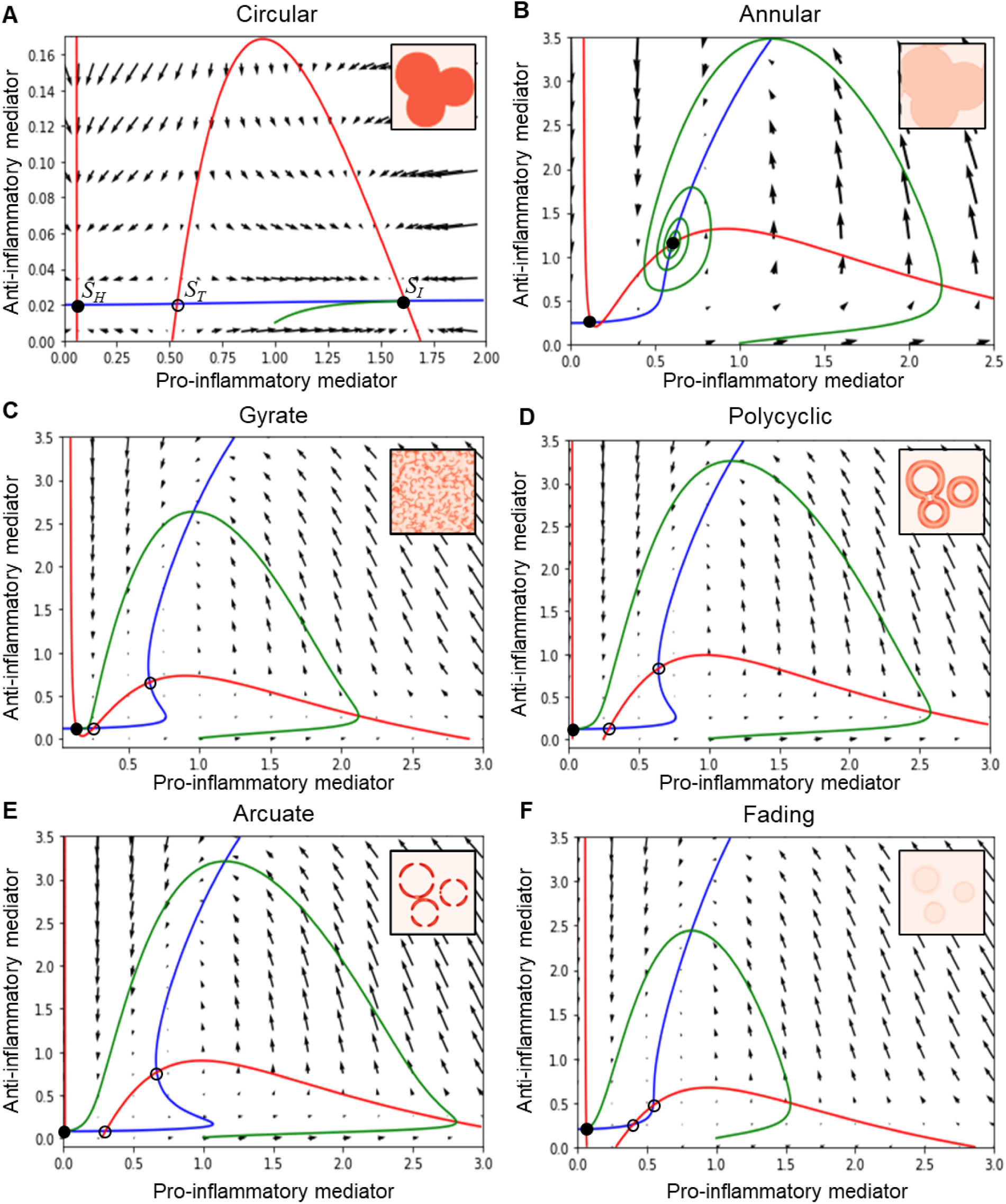
Dynamical characters underlying the five expanding and fading pattern types. The phase space of pro- and anti-inflammatory mediator concentrations depicts the time course (green curve) upon stimulation and the nullclines (red curve for *da/dt* = 0 in Eq. 2a; blue curve for *di/dt* = 0 in Eq. 2b; *D_a_* = *D_i_* = 0). The intersections of the nullclines indicate steady states, where filled and hollow circles represent stable and unstable states, respectively. The time course shows convergence to a stable steady state. Vector fields are also shown to represent mediator dynamics at the respective concentration. The parameter values for each simulation are listed in Table S2.

Bistability also existed in the annular pattern (Fig. 5B). For the circular and annular patterns, the concentrations of pro- and anti-inflammatory mediators eventually reached the inflamed state (Fig. 5A and B). In contrast, in the gyrate, polycyclic, arcuate, and fading patterns, the regulatory feedback resulted in one stable and two unstable steady states, and the mediator concentrations eventually reached a healthy state (Fig. 5C–F). A major difference from the bistability exhibiting circular and annular patterns is the excitability, where the inflamed state is no more stable thereby only appearing in a transient manner upon stimulation. While the excitability and bistability underlie the fading and circular/annular patterns, respectively, consistently with the previous study [Valeyev et al. 2010], the present results further show that the excitability underlies some of the chronic inflammation resulting in gyrate, polycyclic, and arcuate patterns as well.

The distance between the healthy state (*S_H_*) and the threshold state (*S_T_*, a closer unstable steady state to *S_H_*) represents the degree of stability of the healthy against stimulations. We found that the distance was the smallest in the gyrate pattern and increased in the order of polycyclic, arcuate, and fading patterns (Fig. 5C–F). These results indicate that the degree of stability of the healthy state, as well as the presence or absence of stability of the inflamed state, differs between erythema patterns. Therefore, erythema patterns on the skin surface reflect the dynamic balance in the stability of the healthy and inflamed states within the skin.

## Discussion

### Diffusive mediator feedback spatiotemporally regulates erythema patterns between healthy and diseased skin

The spatiotemporal regulation of inflammation is an important theme in biomedical research. Inflammation depends on the feedback of pro- and anti-inflammatory mediators; however, it remains unclear how the feedback regulates fading erythema in healthy skin and expanding erythema in diseased skin. Here, a reaction-diffusion model with mediator feedback (Fig. 1) successfully reproduced the fading patterns (Fig. 2) and five types of expanding patterns (Fig. 3), suggesting that feedback and diffusion can generate fading patterns in healthy skin and expanding patterns in eleven diseases (Table 1). The present study showed that alterations in mediator production destabilized a stable steady state representing a healthy condition while in turn stabilizing the inflamed state (i.e., bifurcation from excitability to bistability [Strogatz 2015]) and led to a transition from a fading pattern to five types of expanding patterns (Fig. 5). In conclusion, the feedback dynamics and diffusion of pro- and anti-inflammatory mediators commonly recapitulated erythema patterns in healthy and multiple diseased skins.

### Prospective treatment from the model prediction

Mediator feedback parameter-dependent transitions from each expanding to fading pattern (Fig. 3, 4, and S2 Fig. A) suggest effective treatment strategies depending on skin barrier conditions. Experimental findings demonstrated that the maximum production rate (*Q_A_* in Eq. 1a and *q_a_* in Eq. 2a) and basal secretion rate (*P_A_* in Eq. 1b and *p_a_* in Eq. 2b) of pro-inflammatory mediators are significantly lower in healthy skin than in diseased skin with a deterioration of the skin microbiome [Pasparakis et al. 2014, Belkaid and Harrison 2017, Meisel et al. 2018] and in diseased skin with defects in the integrity of physical barriers, respectively [Otani and Furuse 2020, Bäsler and Brandner 2017].

Observation of erythema patterns under different skin barrier conditions reveals the influence of skin barrier conditions on the model parameters and thus provides potential treatments to reduce the maximum production rate or the basal secretion rate of pro-inflammatory mediators. For example, probiotics, which improve the composition of the skin microbiome, significantly reduce the maximum production rate of pro-inflammatory mediators [Chen et al. 2020, Yu et al. 2020]. Additionally, probiotics can improve the integrity of physical barriers [Chen et al. 2020], thus reducing the basal secretion rate of pro-inflammatory mediators. Therefore, probiotics can be a prospective treatment, leading to a fading pattern. Further experimental studies on the influence of skin barrier conditions on erythema patterns will offer deeper insights into the development of effective treatments for erythema associated with inflammatory skin disease.

### Applicability of the present model

This study provides a systematic definition of disease severity using this model. The model describes the expanding patterns and fading patterns on the same parameter space (Fig. 4C), which represents how far each expanding pattern is from the fading pattern. This distance is similar to the state-space representation of inflammatory responses, where disease severity is measured as the distance between a patient’s coordinates and that of one of the disease states [Rixen et al. 1996, Voit et al. 2009]. Defining disease severity as the distance between the fading pattern and erythema patterns on the patient’s skin will help estimate the appropriate dosage and strength of treatment for each patient based on their erythema pattern.

Our framework can also predict the disease risk in healthy individuals. The model showed that the fading patterns disappeared at different speeds, in accordance with the parameters (Fig. 2C and D). This means that the parameters in healthy individuals can be estimated by measuring fading speeds using patch tests. Utilizing the obtained parameters, the disease risk of each individual can be evaluated as the distance from the parameter that shows the expanding patterns. Therefore, we propose that the future integration of models, experimental findings, and clinical data will allow for the development of personalized treatment and prediction of inflammatory skin diseases in a non-invasive manner.

### Future implications

The expanding patterns continued to expand in the present model simulations (Fig. 3), while the actual erythema typically stopped expanding and maintained its size in the clinical observation [Shimizu 2017, Seirin-Lee et al. 2020]. This is probably because the present model focuses on the innate immune response, whereas the skin initiates an acquired immune response in the persistence of the innate immune response. Therefore, incorporating the acquired immune response into the model will reproduce the end of the expansion.

Additionally, we focused on well-circumscribed erythema with clear boundaries (Fig. 3) that resulted from inflammation in the upper layers of the skin. Inflammation in the deeper layers of the skin leads to poorly circumscribed erythema with a gradual transition between the affected area and healthy skin [Iryojohokagaku-kenkyusho 2020]. Future studies incorporating the three-dimensional structure of the skin into the present model would take into account poorly circumscribed erythema.

## Conclusions

Here, positive and negative feedback and diffusion of pro- and anti-inflammatory mediators were demonstrated to commonly account for the fading patterns in healthy skin and five types of expanding patterns in diseased skin. Mechanistically, alterations in mediator production destabilize a healthy state while stabilizing an inflamed state, resulting in a transition to diverse expanding patterns. The mediator feedback dynamics is the fundamental regulator of the diseased-to-healthy transition, suggesting effective treatment strategies for each expansion pattern. Therefore, regulating mediator production provides an experimentally testable framework for the spatiotemporal regulation of erythema, which can facilitate the development of a noninvasive and personalized treatment for inflammatory skin diseases.

## Methods

### Source of information on the erythema patterns

First, clinical reports of erythema in the literature were reviewed to examine the association between erythema patterns and skin diseases. Expanding patterns were observed in eleven different diseases, including psoriasis, lupus erythematosus, bullous pemphigoid, Lyme disease, erythema multiforme, lymphoma, annular erythema, Sjögren’s syndrome, sweet syndrome, nummular eczema, and erythema gyratum repens [Tsuji 2002]. We collected clinical photographs of erythema observed in patients whose photographs were extracted from clinical studies using literature searches in PubMed. For example, photographs of psoriasis have been reviewed using the “(psoriasis AND clinical AND pattern) OR (psoriasis AND clinical AND shapes) OR (psoriasis AND clinical spectrum)” search phrases. After reviewing the titles and abstracts, 132 relevant papers with clinical photographs were selected (Table S1).

### Development of the reaction-diffusion model

A reaction-diffusion model was developed to investigate whether regulatory feedback and diffusion of pro- and anti-inflammatory mediators can generate erythema patterns. As pro-inflammatory mediators induce erythema through vasodilation, we used the concentration of pro-inflammatory mediators as an indicator of erythema. The variables of the model reflect the concentrations of pro-inflammatory mediators (*A*) and anti-inflammatory mediators (*I*). Pro-inflammatory mediators are present at low levels in the unstimulated skin through basal secretion [Bäsler and Brandner 2017]. In response to stimulation, keratinocytes and immune cells in the skin secrete pro-inflammatory mediators, which induce their production through positive feedback [Bonizzi and Karin 2004, Nestle et al. 2009]. Pro-inflammatory mediators also induce the production of anti-inflammatory mediators through negative feedback [Opal and DePalo 2000, Zhang and An 2007]. The positive and negative feedback between pro- and anti-inflammatory mediators is shown schematically in Fig. 1I. The production rate of pro-inflammatory mediators is biologically limited; therefore, the model function of *A* saturates the Hill function with the Hill coefficient representing cooperativity in the regulation, *n*. Pro-inflammatory mediators are assumed to degrade naturally at a constant rate [Zhao 2005]. To model these processes, the production of pro-inflammatory mediators (*A*) is represented by the autoregulation of *A* and repression by *I*:

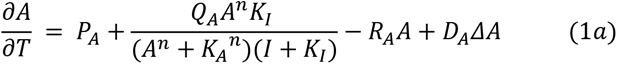

The first term *P_A_* represents the basal production rate. The term 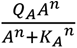 captures the positive feedback, where *Q_A_* and *K_A_* are the maximum production rate and threshold of production of pro-inflammatory mediators, respectively. The term 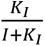 modulates the inhibitory effects of anti-inflammatory mediators. The third and fourth terms represent the degradation with *R_A_* and diffusion *D_A_* with *Δ* denoting the Laplacian operator 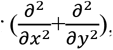, respectively.

Anti-inflammatory mediators induce their production through positive feedback [Zhang and An 2007, Chuang et al. 2016]. Anti-inflammatory mediators, such as pro-inflammatory mediators, are assumed to be present at low levels in the skin through basal secretion and naturally degrade at a constant rate. To model these processes, the production of anti-inflammatory mediators (*I*) was modeled as follows:

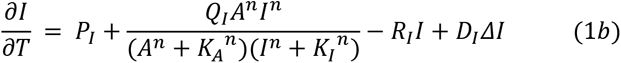

The first, third, and fourth terms in Eq. 1b represent the basal secretion, degradation, and diffusion of anti-inflammatory mediators at *P_I_*, *R_I_*, and *D_I_*, respectively. The second term of Eq. 1b represents the induction of anti-inflammatory mediators by pro-inflammatory mediators and via the positive feedback of anti-inflammatory mediators, where *Q_I_* denotes the maximum production rate of anti-inflammatory mediators.

The values of these parameters depend on the skin conditions. For example, experiments have suggested that the maximum production rate (*Q_A_*) of one type of pro-inflammatory mediator, IL-1β, increases with the deterioration of the skin microbiome [Meisel et al. 2018] and that the basal secretion rate (*P_A_*) of IL-1β increases with a defect in skin barrier integrity [Bäsler and Brandner 2017]. Because of the lack of sufficient quantitative information on the kinetic parameter values and diffusion coefficients, we investigated the model dynamics for a wide range of parameters. For this purpose, the model was nondimensionalized using the following scaling:

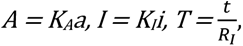

where time is scaled with the degradation rate of anti-inflammatory mediators, which is expected to be in the order of minutes [Baker et al. 2017].

The final system of partial differential equations for pro- and anti-inflammatory mediators is given by:

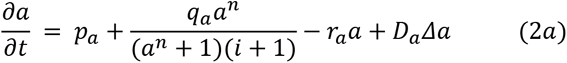

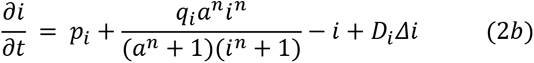

where 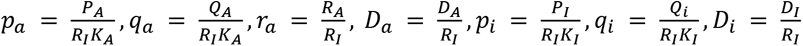.

The meanings of these new parameters are summarized in Table 2. We set the Hill coefficient *n* to 2 to introduce the simplest form of cooperativity. Hence, a reaction-diffusion equation consisting of pro- and anti-inflammatory mediators was used to simulate the development of erythema patterns.

### Numerical simulation of the model

The development of the erythema pattern was simulated by numerically solving the initial value problem in Eq. 2 using the classic Runge-Kutta method: The simulation was performed for cells aligned in a two-dimensional geometry with a periodic boundary condition. As an initial condition of the simulation, pro- and anti-inflammatory mediators were uniformly set as 0.01 in the entire space (Fig. 2, time = 0). Stimulation was introduced into these cells to induce erythema. For stimulation, we referred to the physiological condition at the onset of erythema, where a few small (∼1 mm) inflamed areas exhibited a high concentration of pro-inflammatory mediators [Shimizu 2017]. Accordingly, for each inflamed area, we set a circular area with a high concentration of pro-inflammatory mediators, given by a two-dimensional Gaussian distribution after some time steps (Fig. 2C and D, time = 1). Given the initial conditions and stimulations, we investigated whether the reaction-diffusion model could reproduce erythema patterns.

## Acknowledgments

We would like to show our appreciation to Prof. M. Ueda and Prof. M. Okada (Osaka Univ., Japan) for their insightful suggestions. We also would like to express our gratitude to Dr. M. S. Kitazawa and Dr. K. Matsushita for the stimulating discussions.

## Data availability statement

All relevant data are within the paper and its Supporting Information files.

## Supporting information captions

**Table S1. List of references for erythema observed in the eleven diseases. Table S2. Parameter values used in the simulations.** The parameter values used to generate the fading patterns in Fig. 2 (**A**) and the five types of expanding patterns in Fig. 3 (**B**).

**S1 Fig. Temporal evolution of mediator concentrations in the fading or expanding patterns.** The red and blue lines represent the concentrations of the pro- and anti-inflammatory mediators, respectively. *D_a_* = *D_i_* = 0; and the other parameter values for these simulations are listed in Table S2. **S2 Fig. Pattern selection in the parameter space of pro- and anti-inflammatory mediator productions.**

Fading (F), arcuate (Ar), polycyclic (P), gyrate (G), annular (An) and circular (C) patterns emerged as the steady state (Eq. 2) at the parameter values of *q_a_* and *r_a_* (**A**), *q_a_* and *q_i_* (**B**), *p_a_* and *p_i_* (**C**), *p_a_* and *q_a_* (**D**).

Simulations of the gray areas did not correspond to any of the five patterns. *p_a_* = 0.02, *p_i_* = 0.12, *q_i_* = 6.0 for (**A**), *p_a_* = 0.02, *r_a_* = 0.8, *p_i_* = 0.12 for (**B**), *q_a_* = 3.0, *r_a_* = 0.8, *q_i_* = 6.0 for (**C**), *r_a_* = 0.8, *p_i_* = 0.12, *q_i_* = 6.0 for (**D**). In all simulations, *D_a_* = *D_i_* = 0.3.

**S3 Fig. Alterations in the production rates of pro- and anti-inflammatory mediators transition from fading patterns to various expanding patterns.**

Representation of the patterns generated using Eq. 2 for different values of the parameters *p_a_* and *r_a_* (**A**), *p_a_* and *q_i_* (**B**), *r_a_* and *p_i_* (**C**), and *p_i_* and *q_i_* (**D**). Simulations of the gray areas did not correspond to any of the five patterns. *q_a_* = 3.0, *p_i_* = 0.12, *q_i_* = 6.0 for (**A**), *q_a_* = 3.0, *r_a_* = 0.95, *p_i_* = 0.12 for (**B**), *p_a_* = 0.05, *q_a_* = 3.0, *q_i_* = 6.0 for (**C**), and *p_a_* = 0.05, *q_a_* = 3.0, *r_a_* = 0.95 for (**D**). In all the simulations, *D_a_* = *D_i_* = 0.3.

