## Supporting Materials for "Diffusive mediator feedback explains the health-to-disease transition of skin inflammation"

| | Circular | Annular | Polycyclic | Arcuate | Gyrate |
|---|---|---|---|---|---|
| Psoriasis | [Christophers 2001], [Meier and Sheth 2009] | [Morris et al. 2001], [Tsuji 2002] | [Morris et al. 2001] | [Buxton 1987] | [Jablonska et al. 2000] |
| Lupus erythematosus | N/A | [Cassis and Callen 2005], [Frances et al. 2016], [Herrero et al. 1988], [Maciejewski 1980], [Trayes et al. 2018], [Weston et al. 1999] | [Clark, Shi et al. 2017] | [Cassis and Callen 2005], [Rémy-Leroux 2008], [Toledo-Alberola and Betlloch-Mas 2010] | [Da Silva Sousa et al. 2019], [Kuhn et al. 2007], [Petty et al. 2020], [Sontheimer 1989] |
| Bullous pemphigoid | [Ogilvie et al. 2000], [Reichel et al. 2020] | [Maciejowska et al. 1987], [Ogilvie et al. 2000], [Raposo et al. 2017] | N/A | [Cohen 2016], [Koga et al. 2017], [Maciejowska et al. 1987] | N/A |
| Lyme disease | [Rebman et al. 2021], [Wormser et al. 2005] | [Rebman et al. 2021], [Wormser et al. 2005] | [Mayer et al. 2018], [Rebman et al. 2021], [Schotthoefer et al. 2022], [Wormser et al. 2005] | N/A | N/A |
| Erythema multiforme | [Amode et al. 2018], [Charlesworth 1996], [Lerch et al. 2018] | N/A | [Assier et al. 1995], [Atzori et al. 2003], [Bastuji-Garin et al. 1993], [Cook et al. 2022], [Côté et al. 1995], [Paulino et al. 2018], [Senger et al 2021], | [Drago et al. 1995] | N/A |

| | | | | | |
|---|---|---|---|---|---|
| | | | [Trayes et al. 2018] | | |
| Lymphoma | [Tomasini et al. 2017], [Yumeen and Girardi 2020] | N/A | N/A | [Guitart et al. 2012], [Palacios-Álvarez et al. 2015] | N/A |
| Annular erythema | N/A | [Lee 2000], [Toledo-Alberola and Betlloch-Mas 2010], [Tsuji 2002], [Ziemer et al. 2009] | N/A | [Mu et al. 2015], [Ziemer et al. 2009] | N/A |
| Sjögren syndrome | N/A | N/A | N/A | [Fukunaga et al. 2017], [Tsukazaki et al. 2002] | [Kuhn et al. 2000] |
| Sweet's syndrome | [Gropper 2001], [Clark, Sarcon et al. 2017] | N/A | N/A | N/A | N/A |
| Nummular eczema | [Martínez-Blanco et al. 2016], [Trayes et al. 2018] | N/A | N/A | N/A | N/A |
| Erythema gyratum repens | N/A | N/A | N/A | N/A | [Clyatt and Cole 2022] |

**Table S1. List of references for erythema observed in the eleven diseases.**

| (**A**) Parameter | Linear, Reticular, Circular ($p_i$=0.2) | Circular ($p_i$=0.08) |
|---|---|---|
| $p_a$ | 0.05 | 0.05 |
| $q_a$ | 3 | 3 |
| $r_a$ | 0.95 | 0.95 |
| $p_i$ | 0.2 | 0.08 |
| $q_i$ | 6.5 | 6.5 |
| *Maxstep* | 30 | 30 |

| (**B**) Parameter | Circular | Annular | Polycyclic | Arcuate | Gyrate |
|---|---|---|---|---|---|
| $p_a$ | 0.05 | 0.05 | 0.02 | 0.01 | 0.08 |
| $q_a$ | 2.1 | 3 | 3.1 | 3 | 3 |
| $r_a$ | 0.95 | 0.7 | 0.8 | 0.8 | 0.95 |
| $p_i$ | 0.02 | 0.24 | 0.12 | 0.08 | 0.12 |
| $q_i$ | 6 | 6 | 6 | 6 | 6 |
| *Maxstep* | 300 | 100 | 80 | 50 | 500 |

**Table S2. Parameter values used in the simulations.**

The fading patterns in Fig. 2 (**A**) and the five types of expanding patterns in Fig. 3 (**B**). $D_a = D_i = 0.3$ for all the cases. The simulations run for a total time of *Maxstep*.

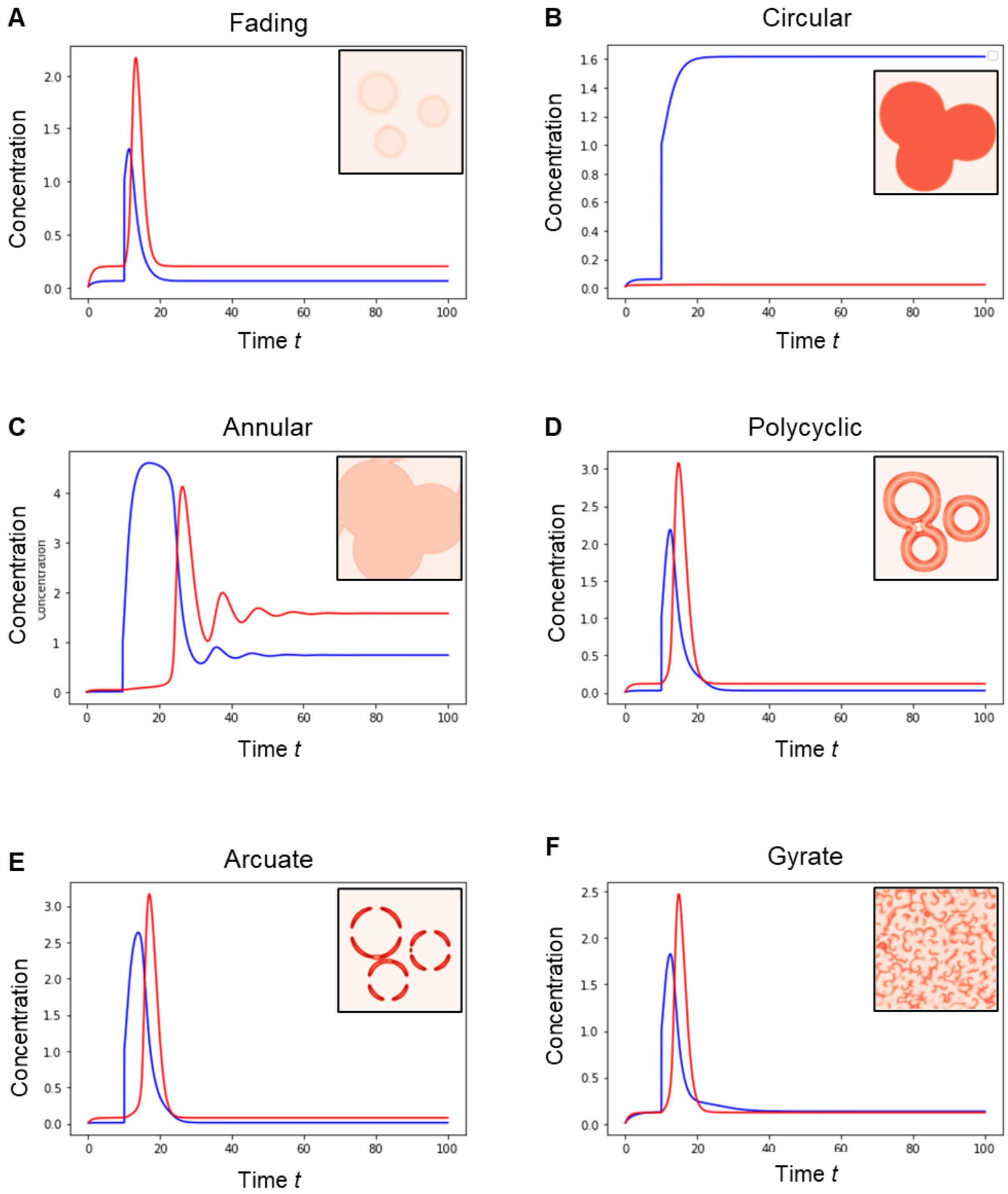


**S1Fig. Temporal evolution of mediator concentrations in the fading or expanding patterns.**

The red and blue lines represent the concentrations of the pro- and anti-inflammatory mediators, respectively. $D_a$=$D_i$=0; and the other parameter values for these simulations are listed in Table S2.

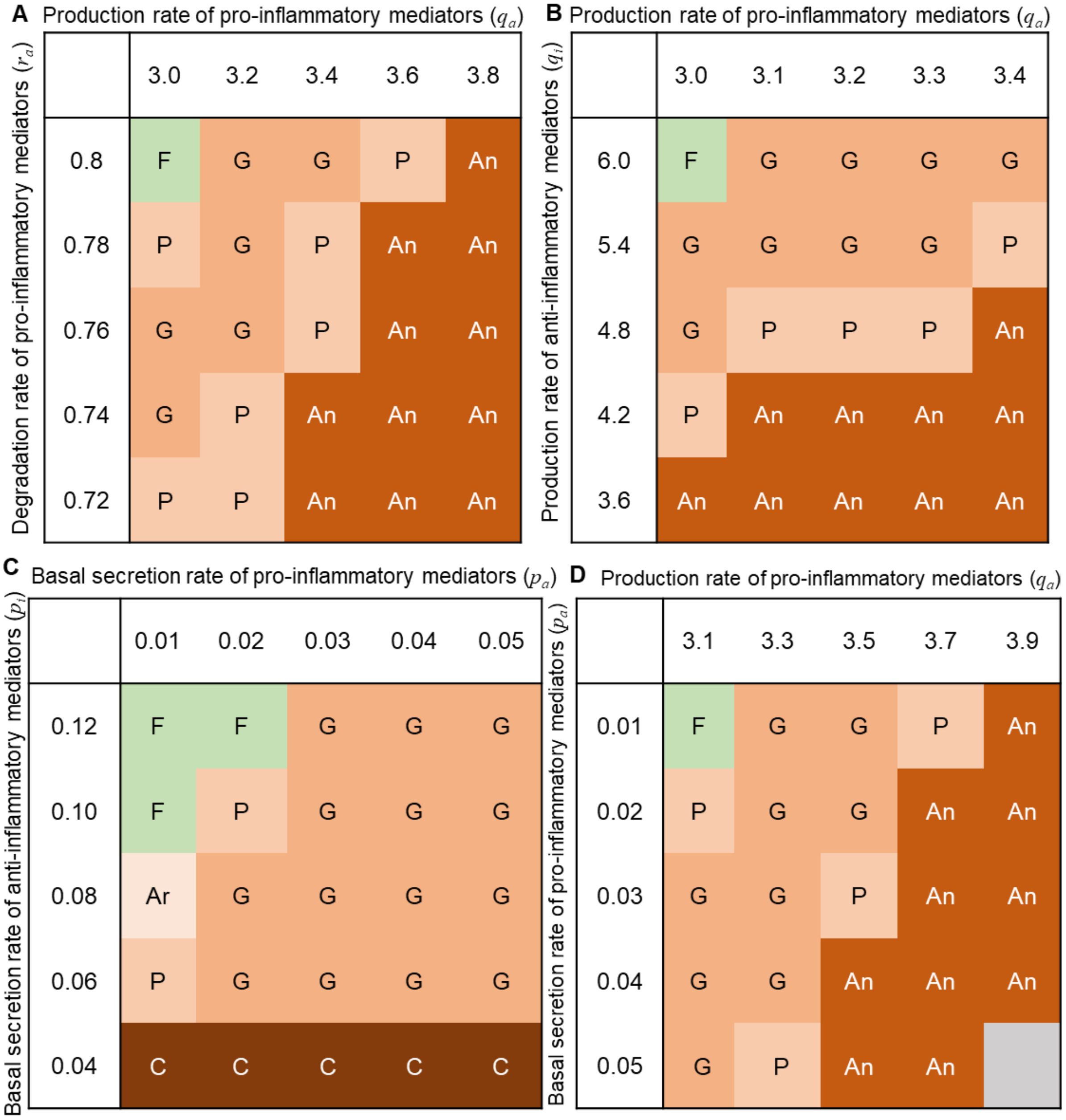


**S2Fig. Pattern selection in the parameter space of pro- and anti-inflammatory mediator productions.**

Fading (F), circular (C), annular (An), polycyclic (P), arcuate (Ar), and gyrate (G) patterns emerged as the steady state (Eq. 2) at the parameter values of $q_a$ and $r_a$ (**A**), $r_a$ and $q_i$ (**B**), $p_a$ and $p_i$ (**C**), $p_a$ and $q_a$ (**D**). Simulations of the gray areas did not correspond to any of the five patterns. $p_a$=0.02, $p_i$=0.12, $q_i$=6.0 for

(**A**), $p_a$=0.02, $r_a$=0.8, $p_i$=0.12 for (**B**), $q_a$=3.0, $r_a$=0.8, $q_i$=6.0 for (**C**), $r_a$=0.8, $p_i$=0.12, $q_i$=6.0 for (**D**). In all simulations, $D_a$=$D_i$=0.3.

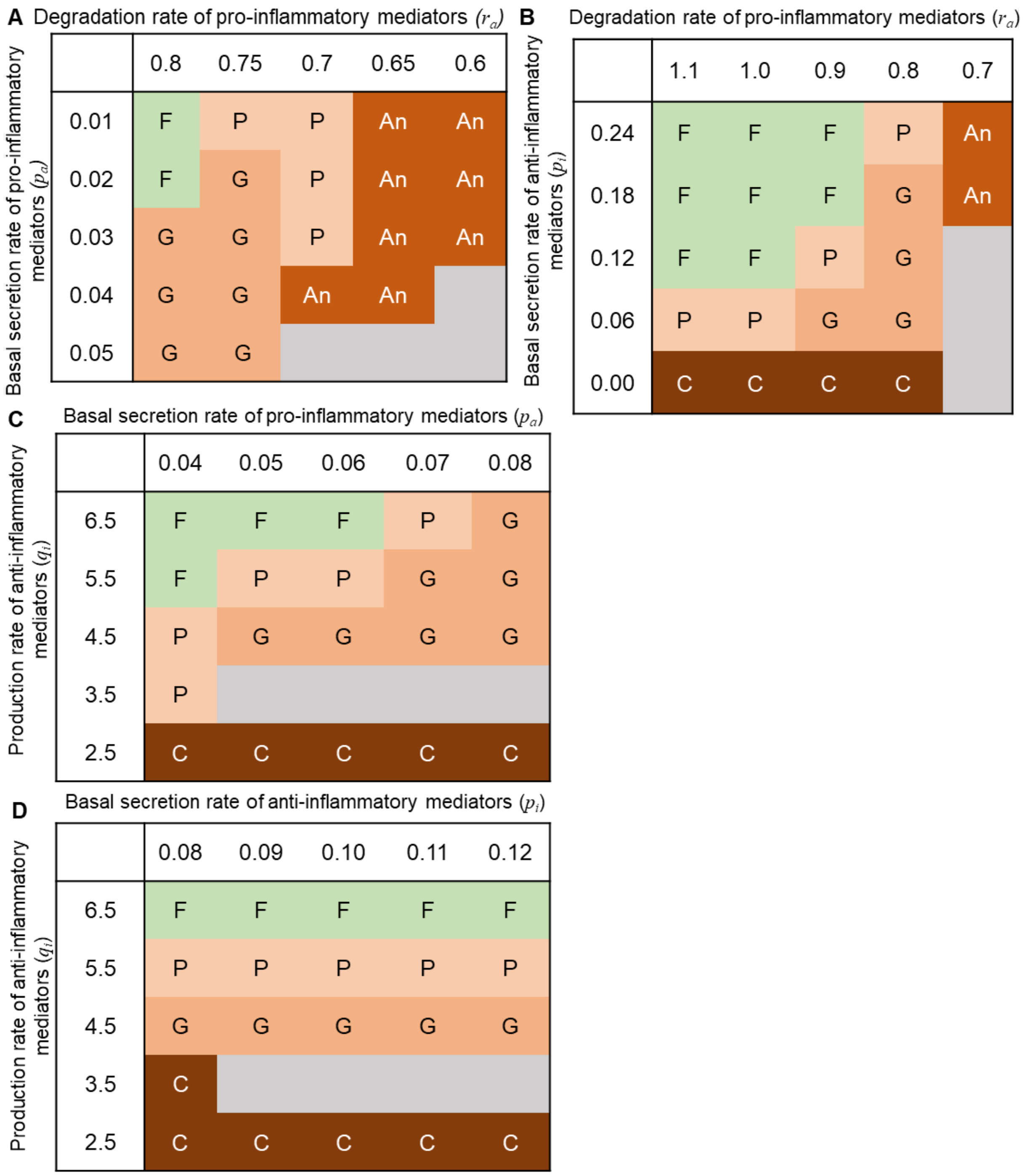


**S3Fig. Alterations in the production rates of pro- and anti-inflammatory mediators transition from fading patterns to various expanding patterns.**

Representation of the patterns generated using Eq. 2 for different values of the parameters $p_a$ and $r_a$ (**A**), $p_a$ and $q_i$ (**B**), $r_a$ and $p_i$ (**C**), and $p_i$ and $q_i$ (**D**). Simulations of the gray areas did not correspond to any of the five patterns. $q_a$=3.0, $p_i$=0.12, $q_i$=6.0 for (**A**), $q_a$=3.0, $r_a$=0.95, $p_i$=0.12 for (**B**), $p_a$=0.05, $q_a$=3.0, $q_i$=6.0 for (**C**), and $p_a$=0.05, $q_a$=3.0, $r_a$=0.95 for (**D**). In all the simulations, $D_a$=$D_i$=0.3.
